# Ablation of sympathetic nerve-β3 adrenergic receptor-mediated adipose tissue lipolysis attenuates alcohol-induced liver injury in mice

**DOI:** 10.1101/2024.12.08.627372

**Authors:** Piumi Wickramasinghe, Alexandre Caron, Sreeja Eadha, Preethi Parupalli, Sarvani Ganapavarapu, Joel Elmquist, Chen Liu, Lin Jia

## Abstract

**BACKGROUND & AIMS:** Binge drinking causes fat accumulation in the liver and is a known risk factor for more severe forms of alcohol-associated liver disease (ALD). Although adipocyte-released free fatty acids (FFA) have been shown to contribute to alcohol-induced liver damage, the signaling pathways that trigger lipolytic activity in adipose tissues following acute alcohol overconsumption is largely unknown. Notably, activation of sympathetic nerve-β3 adrenergic receptor (ADRB3) plays a central role in sustained adipocyte lipolysis. However, whether this pathway is involved in acute alcohol-induced lipolysis remains unclear. We aimed to explore the effect of the sympathetic nerve-ADRB3-mediated pathway on adipocyte lipolytic action and fatty liver development following acute alcohol exposure.

**METHODS:** C57BL/6J mice were administered a single binge of alcohol to model acute alcohol exposure. 6-hydroxydopamine (6-OHDA) was injected systemically or locally to ablate sympathetic nerves. Mice lacking Adrb3 selectively in fat tissues (Adrb3^FKO^) were generated. White adipose tissue lipolysis, fatty liver development, and liver damage were investigated.

**RESULTS:** A single alcohol binge in C57BL/6J mice led to significant increases in white adipose tissue (WAT) norepinephrine (NE) content and plasma FFA levels, accompanied by the development of alcoholic hepatic steatosis. Acute alcohol-induced adipose tissue lipolysis and ALD were significantly mitigated by 6-OHDA-mediated systemic and fat tissue specific sympathetic nerve ablation. Deletion of Adrb3 in adipocytes protected mice from acute alcohol-induced adipose tissue lipolysis, hepatic fat accumulation, and liver injury.

**CONCLUSION:** Our data indicate that binge drinking leads to the development of fatty liver and liver damage by activating adipose tissue sympathetic nerve-ADRB3-mediated lipolysis in mice.

**SUMMARY:** Binge drinking causes hepatic steatosis and liver injury through the activation of sympathetic nerve-β3 adrenergic receptor-stimulated white adipose tissue lipolysis and release of free fatty acids.

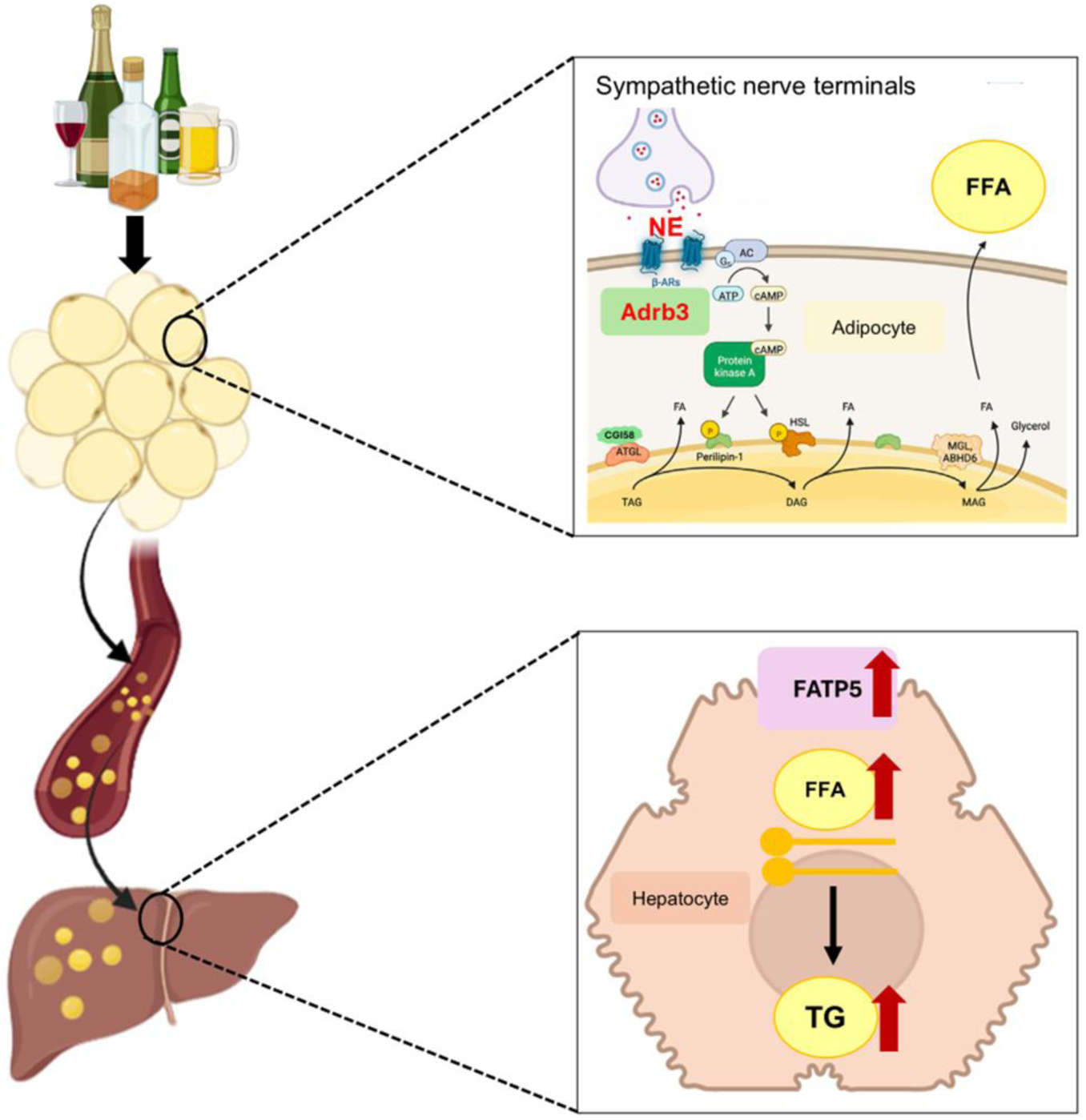

## Introduction

Alcohol-associated liver disease (ALD) ranks among the most prevalent liver diseases globally.^1^ ALD progresses through various stages, starting from steatosis which is characterized by lipid droplet accumulation in hepatocytes. The prevalence of acute alcohol consumption has been rising, with over 58 million Americans reporting binge drinking on at least one occasion within 30 days as reported by The National Survey on Drug Use and Health (NSDUH) in 2010. Importantly, binge drinking not only induces fatty liver but has been considered a significant risk factor for advanced alcoholic liver damage, which can lead to more severe forms of ALD.^2^ Acute/binge drinking is defined by consuming 4 or more drinks for women or consuming 5 or more drinks for men in about two hours.^3^ Acute alcohol exposure can also include a period of heavy drinking that may span several days or periods of intermittent, repeated episodes of heavy drinking. For example, in the clinical setting, an episode of “binge” drinking often describes a period of alcohol consumption and intoxication that lasts upward of 2 days.^4^ Although alcohol-induced organ damage has been studied for decades, the underlying mechanisms of binge drinking-associated ALD remains largely unknown.^2^

Adipose tissues store excess energy as triglycerides in lipid droplets. Under energy-demanding states, these triglycerides are mobilized by a hydrolytic process called lipolysis. Catecholamines are one of the primary regulators of adipocyte lipolysis by binding to the G protein-coupled α and β-adrenergic receptors (β-ARs).^5,6^ The α2 adrenergic receptor couples with Gi-proteins to suppress lipolysis, while the β-ARs interact with Gs-proteins to enhance lipolysis. Among the three β-ARs, β3 adrenergic receptor (Adrb3) is the predominant β-ARs that mediates significant and sustained lipolytic signaling in rodents^6^.

The stimulation of β-ARs happens with the release of norepinephrine (NE) from the sympathetic nervous system (SNS). The binding of NE to Adrb3 activates cAMP-dependent protein kinase (PKA). The subsequent phosphorylation of perilipin 1 allows the interaction between comparative gene identification-58 (CGI-58) and adipose triglyceride lipase (ATGL), causing hydrolysis of triacylglycerol to produce diacylglycerol and free fatty acid (FFA). Phosphorylation of hormone sensitive lipase (HSL) by PKA breaks down diacylglycerol into monoacylglycerol and FFA. Hydrolysis of monoacylglycerols is carried out by monoacylglycerol lipase. Glycerol and FFAs can then be released into the bloodstream for use as an energy source by other tissues in the body.

Findings from both alcoholic patients and preclinical animal models suggest that alcohol consumption is linked to significantly increased lipolytic activities in adipose tissues.^7,8,9–11^ Specifically, subjects consuming more than 5 drinks in men and 2 drinks in women exhibit notable decreases in the percentage of body fat content and significant increases in plasma alanine aminotransferase (ALT) levels.^10,11^ Rodents fed alcohol chronically or exposed to NIAAA drinking model show increased expression of phosphorylated HSL (pHSL) in WAT, elevated circulating FFA and severe liver injury.^7–9^ However, this correlation has not been thoroughly investigated under binge drinking conditions. Although deficiency of adipocyte ATGL or CGI-58 has been reported to attenuate alcohol induced elevations in lipolytic proteins and the development of hepatic steatosis, the upstream signaling pathways that mediate acute alcohol drinking-induced adipose tissue hyper-lipolysis remain to be determined.^7,12^

Adipose tissues are extensively innervated by sympathetic nerve fibers. Additionally, locally released catecholamines from sympathetic nerves are both necessary and sufficient to initiate lipolysis.^6^ In the current study, we demonstrated that a single alcohol binge in C57BL/6J mice led to significant increases in WAT NE content and plasma FFA levels, accompanied by the development of alcoholic hepatic steatosis. Additionally, acute alcohol-induced adipose tissue lipolysis and ALD were significantly mitigated by 6- OHDA-mediated systemic and gonadal WAT (gWAT)-specific SNS ablation. To investigate the role of adipocyte Adrb3 in acute alcohol-induced lipolysis, we generated mice with targeted deletion of Adrb3 in fat depots (Adrb3^FKO^). Our findings revealed that selective ablation of Adrb3 in adipocytes protected mice from acute alcohol-induced increases in plasma FFA, ALT and hepatic triglyceride accumulation.

## Results

### Acute alcohol drinking increases plasma FFA and hepatic triglyceride contents in mice

Prior studies have reported that people who consume more than 5 drinks per day have a lower percent body fat than control non-drinkers.^10,11^ Similarly, when C57BL/6J mice were treated with a single binge of alcohol (5g/kg body weight (BW), EtOH) by oral gavage, these mice exhibited significant reductions in fat pad mass to BW ratios (**Figure 1A-B**), which was correlated with substantial increases in plasma FFA (**Figure 1C**). In addition, significantly increased liver triglyceride contents and plasma ALT levels were observed in alcohol-gavaged mice (**Figure 1D-E**). Acute alcohol treatment did not affect the liver/BW ratio in mice (**Figure 1F**). Collectively, these data indicated the link between adipose tissue lipolysis and binge drinking-induced liver damage. Considering the extensively innervated sympathetic nerves in the adipose tissue and their roles in lipolysis, we set out to determine the NE content in binge alcohol-treated mice. Intriguingly, we found that acute alcohol drinking resulted in significant increases in local NE levels in mouse gWAT and iWAT (**Figure 1G-H**).

**Figure 1.**
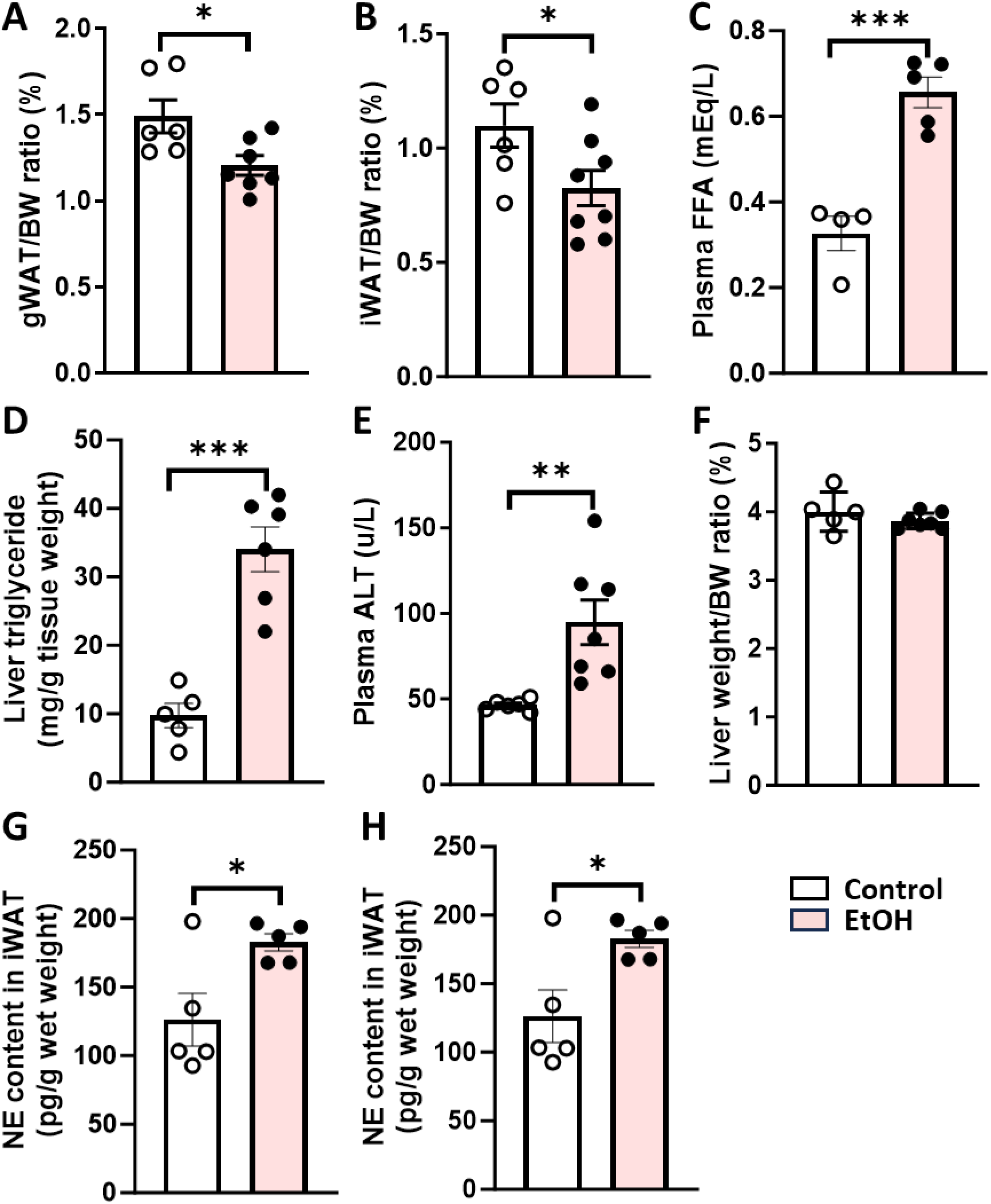
Binge alcohol increases adipose tissue lipolysis, hepatic steatosis and liver damage. C57BL/6J male mice were subjected to a single alcohol binge (EtOH, 5 g/kg BW) or isocaloric maltose dextrin (Control). (A) gWAT/BW ratio (%). (B) iWAT/BW ratio (%). (C) Plasma FFA. (D) Liver triglyceride. (E) Plasma ALT. (F) Liver/BW ratio (%). (G) gWAT NE content. (H) iWAT NE content. n=5-8. Data are expressed as the means ± SEM. *P < .05, **P < .01, *** P < .001.

### Systemic 6-OHDA treatment prevents adipose tissue lipolysis and attenuates fatty liver development after acute alcohol treatment

To determine if SNS function is required for enhanced lipolysis of WAT and the subsequent development of hepatic steatosis after binge alcohol consumption, male C57BL/6J mice underwent chemical sympathectomy by intraperitoneal (ip) injection of 6- hydroxydopamine (6-OHDA), a selective neurotoxin that destroys sympathetic nerves.^13^ This treatment has been shown to effectively block sympathetic nerve activity in various tissues.^13^ We found that a single binge of alcohol led to reductions in gWAT/BW and iWAT/BW ratios in saline-treated mice. In addition, this acute alcohol drinking-induced fat mass decreases were observed in 6-OHDA-exposed iWAT, but not gWAT (**Figure S1A-B**). Prior findings have shown that chronic alcohol drinking is associated with elevated expression of pHSL in adipose tissues.^7,8,12^ We observed that binge drinking caused increased pHSL expression in both gWAT and iWAT as shown in **Figure 2A-B**. Intriguingly, these elevations were dramatically blunted in SNS-ablated mice (**Figure 2A-B**). Although 6-OHDA treatment resulted in increased circulating FFA levels in control-exposed mice, no additional increase was observed in SNS-ablated mice after acute alcohol exposure (**Figure 2C**). In contrast to elevated hepatic triglyceride content in mice following an alcohol binge (**Figure 2D**), 6-OHDA-treated mice exhibited dramatically attenuated triglyceride accumulation in the liver (**Figure 2D**) and significantly reduced liver weight to BW ratio (**Figure S1C**). Alcohol binge caused an elevation in plasma ALT levels (**Figure 2E**). However, similar plasma ALT was observed in control and EtOH- treated mice after SNS ablation (**Figure 2E**). These findings suggested that systemic SNS ablation could partially protect mice from binge drinking-associated adipose tissue lipolysis, fatty liver development and liver damage.

**Figure 2.**
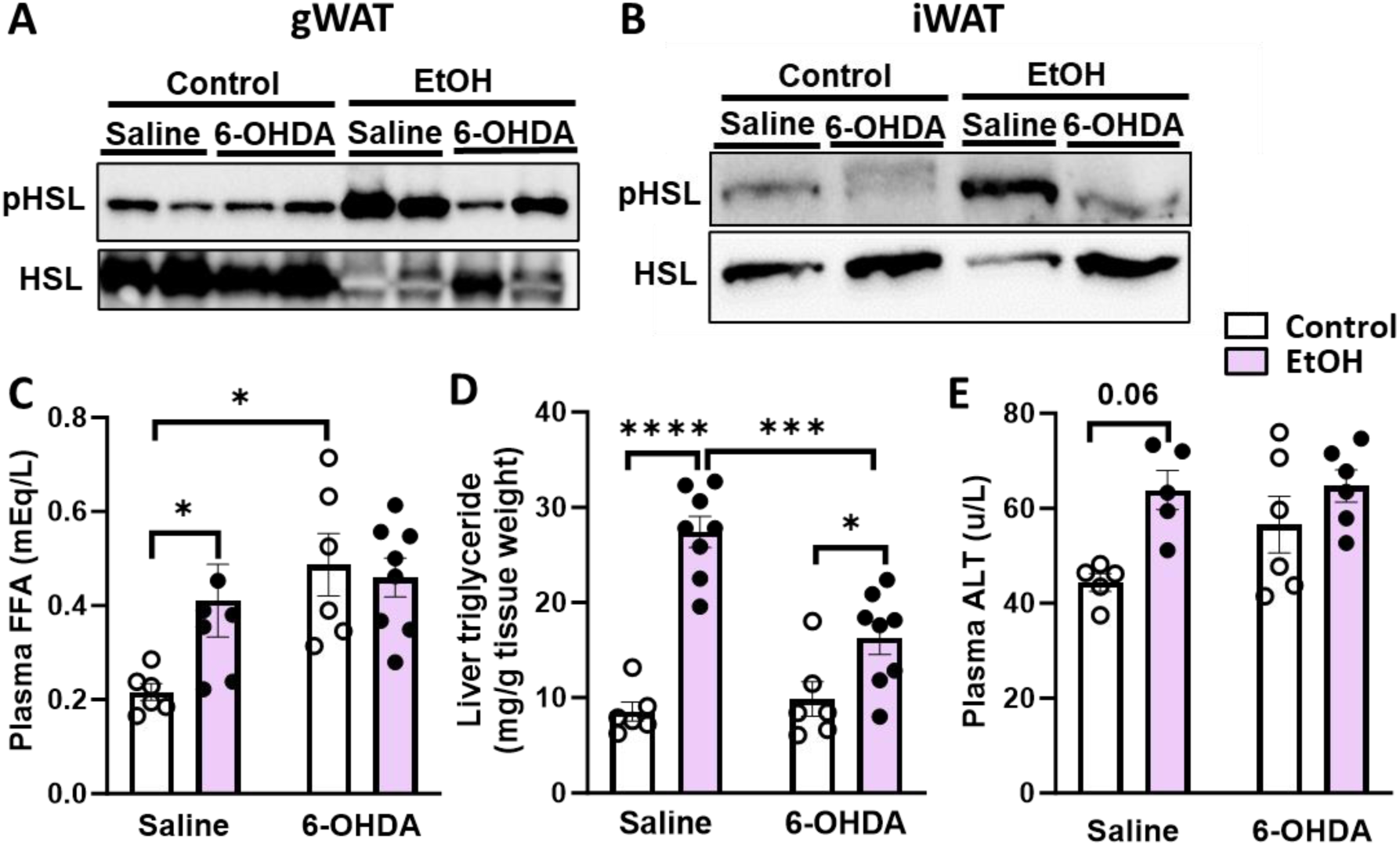
Systemic 6-OHDA treatment attenuates adipose tissue lipolysis and hepatic steatosis. C57BL/6J male mice were intraperitoneally injected with 6-OHDA or vehicle control once per day for 3 days. On the fourth day, mice were subjected to binge drinking and were killed 6 hours after oral gavage. (A-B) pHSL protein levels in (A) gWAT and (B) iWAT. (C) Plasma FFA. (D) Liver triglyceride. (E) Plasma ALT. n=5-8. Data are expressed as the means ± SEM. *P < .05, *** P < .001, **** P < .0001. 6-OHDA, 6-Hyroxydopamine.

### Local injection of 6-OHDA in gWAT attenuates fatty liver development and liver damage following acute alcohol treatment

It has been shown that sympathetic nerve-mediated lipolytic action is more pronounced in gonadal adipocytes than in inguinal adipocytes.^13^ To explore gWAT-mediated lipolysis in acute alcohol induced liver injury, we ablated gWAT sympathetic nerves via local 6- OHDA injection, which has been shown to successfully decrease local NE content.^14^ We observed dramatically attenuated lipolytic function in 6-OHDA-treated gWAT after alcohol treatment, indicated by less pHSL expression (**Figure 3A**). Similarly elevated circulating FFA levels were observed between saline and 6-OHDA-treated mice after alcohol exposure (**Figure 3B**). These could be due to the active uptake of FFA by the livers from saline-injected binge drinking-treated mice. Indeed, these mice displayed a 4-fold increase in hepatic triglyceride content compared to saline-injected, control-exposed mice (**Figure 3C**). Interestingly, binge alcohol-induced hepatic steatosis and plasma ALT elevation were significantly ameliorated by gWAT SNS inactivation (**Figure 3C-D**). These data indicated that sympathetic activity in gWAT partially contributed to acute alcohol-mediated adipose tissue lipolysis, hepatic steatosis, and liver injury.

**Figure 3.**
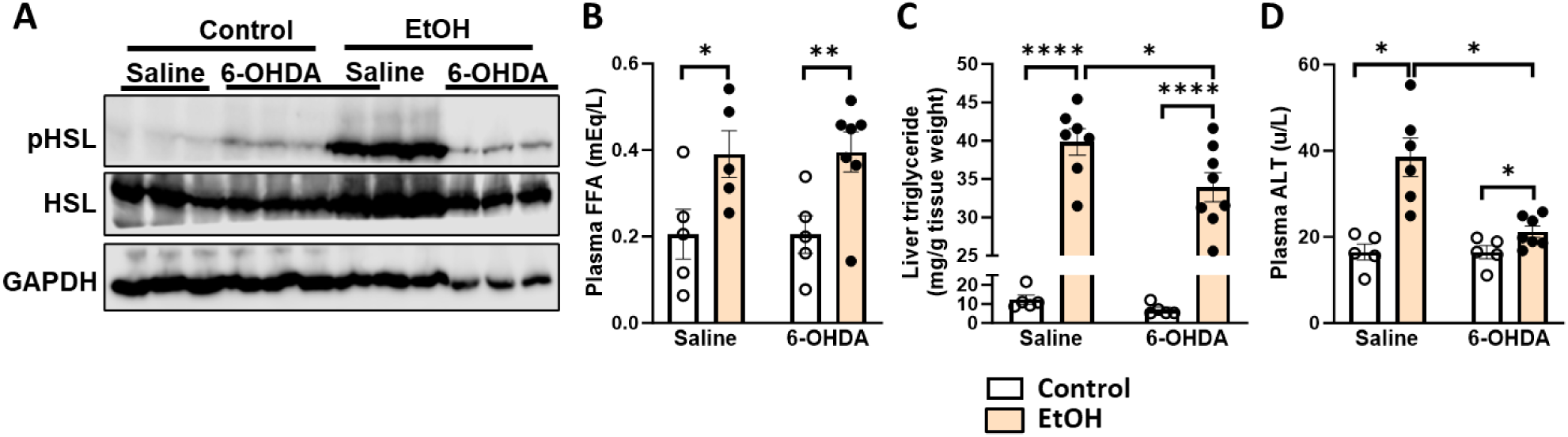
Blocking gWAT sympathetic nerve via local administration of 6-OHDA attenuates adipose tissue lipolysis and fatty liver development. C57BL/6J male mice were subjected to chemical sympathectomy by injecting 6-OHDA into gWAT pads. After two weeks of recovery, mice were given a single alcohol binge. (A) gWAT pHSL protein level. (B) Plasma FFA. (C) Liver triglyceride. (D) Plasma ALT. n=5-8. Data are expressed as the means ± SEM. *P < .05, **P < .01, **** P < .0001. 6-OHDA, 6-Hyroxydopamine.

### Combined treatment of acute alcohol and CL316, 243 exacerbates plasma FFA and liver triglyceride accumulation in mice

NE released from activated sympathetic nerves interacts with either α or β-ARs to regulate adipocyte lipolysis. ADRB3 is the predominant adrenergic receptor in rodents’ adipose tissue which mediates significant and sustained lipolytic signals.^6,15^ To examine the involvement of Adrb3-mediated signaling in acute alcohol-induced hyper-lipolysis, male C57BL/6J mice were oral gavaged with EtOH/control followed by ip injection of saline or CL316,243 (0.1 mg/kg BW), a selective ADRB3 agonist. We found that CL316,243 treatment led to elevated plasma FFA and increased hepatic triglyceride in both control and EtOH-gavaged mice (**Figure 4A-B**). Despite comparable circulating FFA levels between EtOH and EtOH+CL316,243-treated mice 5.5 hours post injections, combined binge drinking and CL316,243 treatment resulted in the most triglyceride accumulation in the liver (**Figure 4B**).

**Figure 4.**
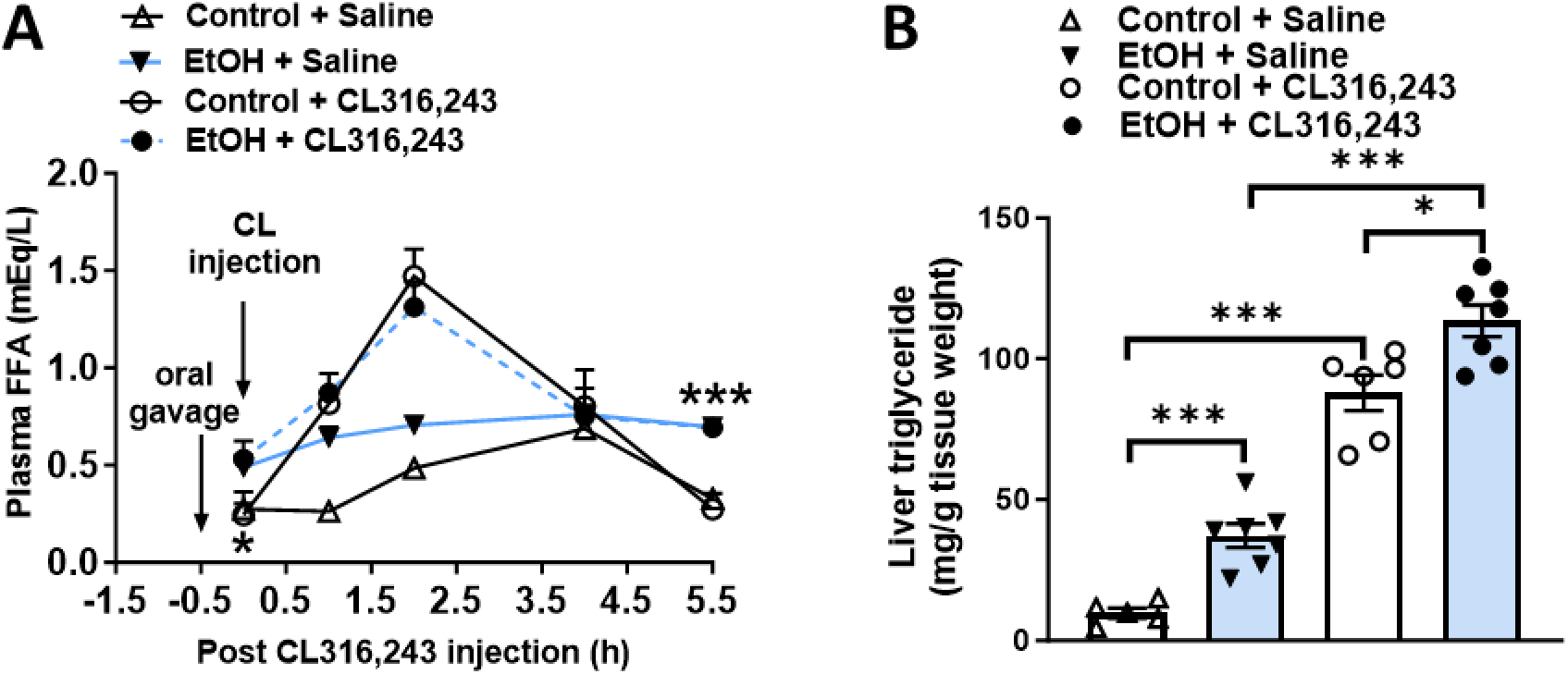
Combination of binge alcohol and CL316,243 treatment exacerbates hepatic steatosis in mice. C57BL/6J male mice were subjected to a single binge of alcohol (EtOH) or isocaloric maltose (Control). 30 minutes later, intraperitoneal injections of saline or CL316,243 were performed and blood was collected at 0, 1, 2, 4, and 5.5 hours after CL316,243 administration. (A) Plasma FFA. (B) Liver triglyceride. n=5-7. Data are expressed as the means ± SEM. *P < .05, *** P < .001.

### Deletion of Adrb3 in adipose tissues dramatically reduces acute alcohol-induced WAT lipolysis

To investigate the role of adipocyte Adrb3 in acute alcohol-induced lipolysis and fatty liver disease, Adrb3^fl/fl^ mice on a C57BL/6J background were generated using CRISPR-Cas9 technology. Two loxP sites were inserted into the upstream of exon 2 and downstream of exon 3 of the Adrb3 DNA sequence (**Figure S2A**) respectively, to allow for conditional Adrb3 deletion. Mice were genotyped for either the wild type (wt) or floxed (fl) allele by PCR using genomic DNA from mouse tails. PCR products of WT and floxed alleles were 274bp and 314bp, respectively (**Figure S2B**). To generate mice lacking Adrb3 in adipocyte (Adrb3^FKO^), Adrb3^fl/fl^ mice were crossed with Adipoq-Cre mouse line. Chow-fed Adrb3^fl/fl^ and Adrb3^FKO^ mice were treated with either saline or CL316,243 (0.1mg/kg BW) via ip injection to validate these mouse models. Consistent with prior reports,^16–18,20^ CL316, 243 treatment reduced blood glucose levels and increased plasma FFA contents in Adrb3^fl/fl^ mice (**Figure 5A-B**). However, these changes were completely blunted in CL316,243-treated Adrb3^FKO^ mice, which were comparable to that of mice treated with saline (**Figure 5A-B**). In addition, dramatically reduced Adrb3 mRNA expressions were found in the gWAT and iWAT of Adrb3^FKO^ mice following either an oral gavage of 5 g/kg BW of EtOH or isocaloric maltose dextrin (control) (**Figure S3A and S3C**). Alcohol feeding did not affect Adrb1 or Adrb2 expression (**Figure S3A**), but significantly suppressed Adra1 and Adra2 mRNA levels in the gWAT of Adrb3^fl/fl^ mice (**Figure S3B**). Adipocyte Adrb3 deficiency was associated with dramatically reduced mRNA expression of Adrb1, Adrb2, Adra1 and Adra2 in gWAT following binge drinking (**Figure S3A-B**). For iWAT, acute alcohol treatment suppressed Adrb1 and increased Adra2 expression (**Figure S3C-D**). Expressions of Adrb2 and Adra1 in iWAT were not influenced by binge drinking or Adrb3 deficiency (**Figure S3C-D**).

**Figure 5.**
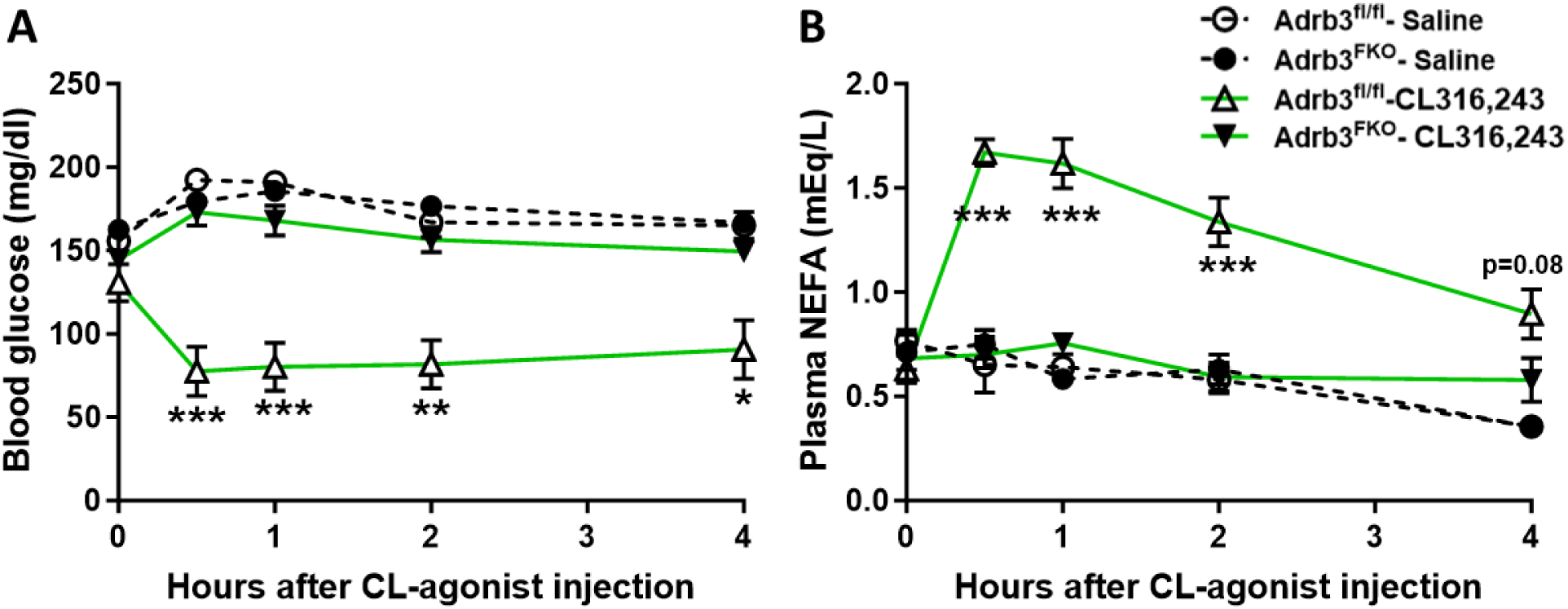
Validation of the Adrb3^FKO^ mouse model. Male Adrb3^FKO^ and their littermate controls (Adrb3^fl/fl^) were treated with saline or CL316,243 (0.1mg/kg BW) via intraperitoneal injection. Blood was collected at 0, 0.5, 1, 2, and 4 hours after CL316,243 administration. (A) Blood glucose. (B) Plasma FFA. n=5-6. Data are expressed as the means ± SEM. *P < .05, **P < .01, *** P < .001.

Then pHSL expression and plasma FFA levels were measured. As expected, alcohol treatment greatly increased pHSL expression in both gWAT and iWAT of Adrb3^fl/fl^ mice (**Figure 6A-B**), which was associated with significantly elevated plasma FFA (**Figure 6C**). However, this acute alcohol-induced hyper-lipolysis was dramatically attenuated in Adrb3^FKO^ mice (**Figure 6A-C**). Hematoxylin and eosin (H&E) stained sections revealed that acute alcohol exposure-induced reductions in adipocyte size in both gWAT and iWAT of Adrb3^fl/fl^ mice were attenuated in Adrb3^FKO^ mice (**Figure 6D-E**). A significant decrease in gWAT/BW ratio was observed in binge drinking-treated Adrb3^fl/fl^, but not Adrb3^FKO^ mice (**Figure S4A**). Regardless of the genotype, alcohol treatment did not affect iWAT/BW ratios (**Figure S4B**).

**Figure 6.**
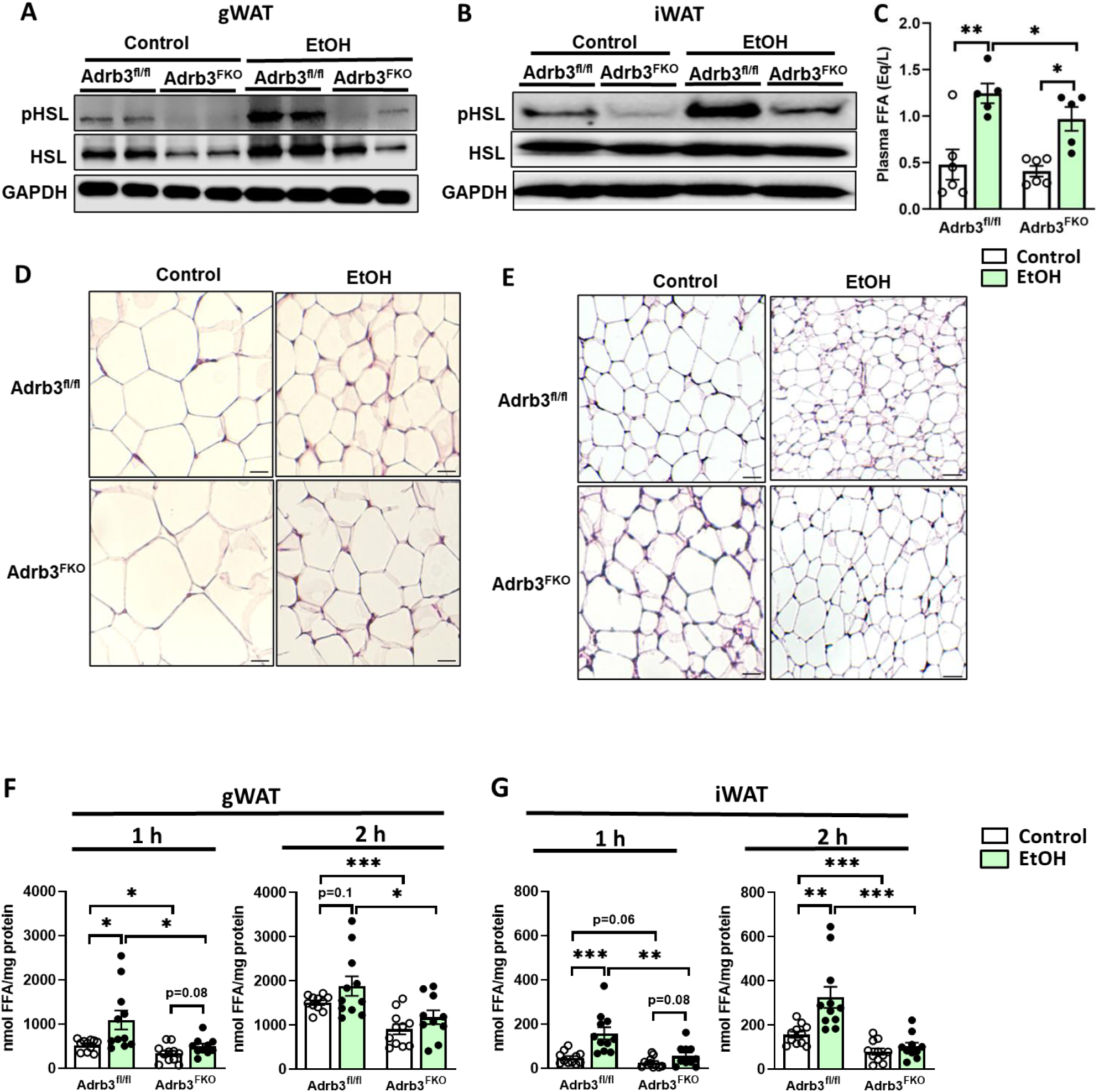
Deletion of Adrb3 in adipose tissues attenuates adipocyte lipolysis in binge drinking-treated mice. Chow-fed Adrb3^fl/fl^ and Adrb3^FKO^ mice were given a single alcohol binge. Mice were killed 6 hours after oral gavage. (A) gWAT pHSL protein expression. (B) iWAT pHSL protein expression. (C) Plasma FFA (n=5-6). (D-E) H&E staining of (D) gWAT and (E) iWAT sections. Scale bar: 50 µm. (F-G) Ex vivo lipolysis assay was performed using adipose tissue explants isolated from Adrb3^fl/fl^ and Adrb3^FKO^ mice subjected to oral gavages of Control or EtOH. FFA contents in the culture medium of (F) gWAT and (G) iWAT. Data are expressed as the means ± SEM. *P < .05, **P < .01, *** P < .001.

Regional variations in adipose tissue lipolysis and release of FFA have been reported.^19^ To study the fat pad-specific role of Adrb3 in lipolysis caused by acute alcohol drinking, *ex vivo* lipolysis assay was performed using gWAT and iWAT isolated from Adrb3^fl/fl^ and Adrb3^FKO^ mice 2 hours after the oral gavage. We found that compared to the control treatment, acute alcohol exposure significantly increased the release of FFA from gWAT and iWAT into the culture media after both 1-hour and 2-hour incubation in Adrb3^fl/fl^ mice (**Figure 6F-G**). This binge drinking-stimulated FFA release was significantly attenuated in gWAT and iWAT of Adrb3^FKO^ mice (**Figure 6F-G**). In addition, Adrb3^FKO^ mice had a lower basal FFA release compared to Adrb3^fl/fl^ mice (**Figure 6F-G**). Relative to iWAT, gWAT showed a much higher capability in FFA release following acute alcohol treatment in both Adrb3^fl/fl^ and Adrb3^FKO^ mice. These results suggested that adipocyte Adrb3 plays a crucial role in driving alcohol-induced lipolysis in both gWAT and iWAT.

### Adipose Adrb3 deficiency dramatically reduces binge alcohol exposure-induced fatty liver development and liver damage

It has been shown that fatty acids resulting from alcohol-induced lipolysis are reverse transported and accumulated in the liver.^21^ Moreover, this adipose tissue-liver crosstalk is associated with increased expression of fatty acid transporter protein 5 (FATP5) in mouse livers following chronic alcohol feeding.^8^ Consistent with this concept, we found that after acute alcohol treatment, hepatic FATP5 expression and liver FFA contents were significantly increased in Adrb3^fl/fl^ mice but greatly blunted in Adrb3^FKO^ mice (**Figure 7A-B**). No alterations were observed for hepatic CD36 expression among the four groups (**Figure S4C**). Dramatically increased hepatic triglyceride and plasma ALT were observed in alcohol-treated Adrb3^fl/fl^ mice (**Figure 7C-D**). In contrast, alcohol-induced fatty liver development and liver damage were partially but significantly blunted in Adrb3^FKO^ mice (**Figure 7C-D**). Furthermore, liver H&E staining displayed elevated lipid droplet accumulation in hepatocytes of acute alcohol-exposed Adrb3^fl/fl^ mice and this elevation was attenuated in Adrb3^FKO^ mice (**Figure 7E**).

**Figure 7.**
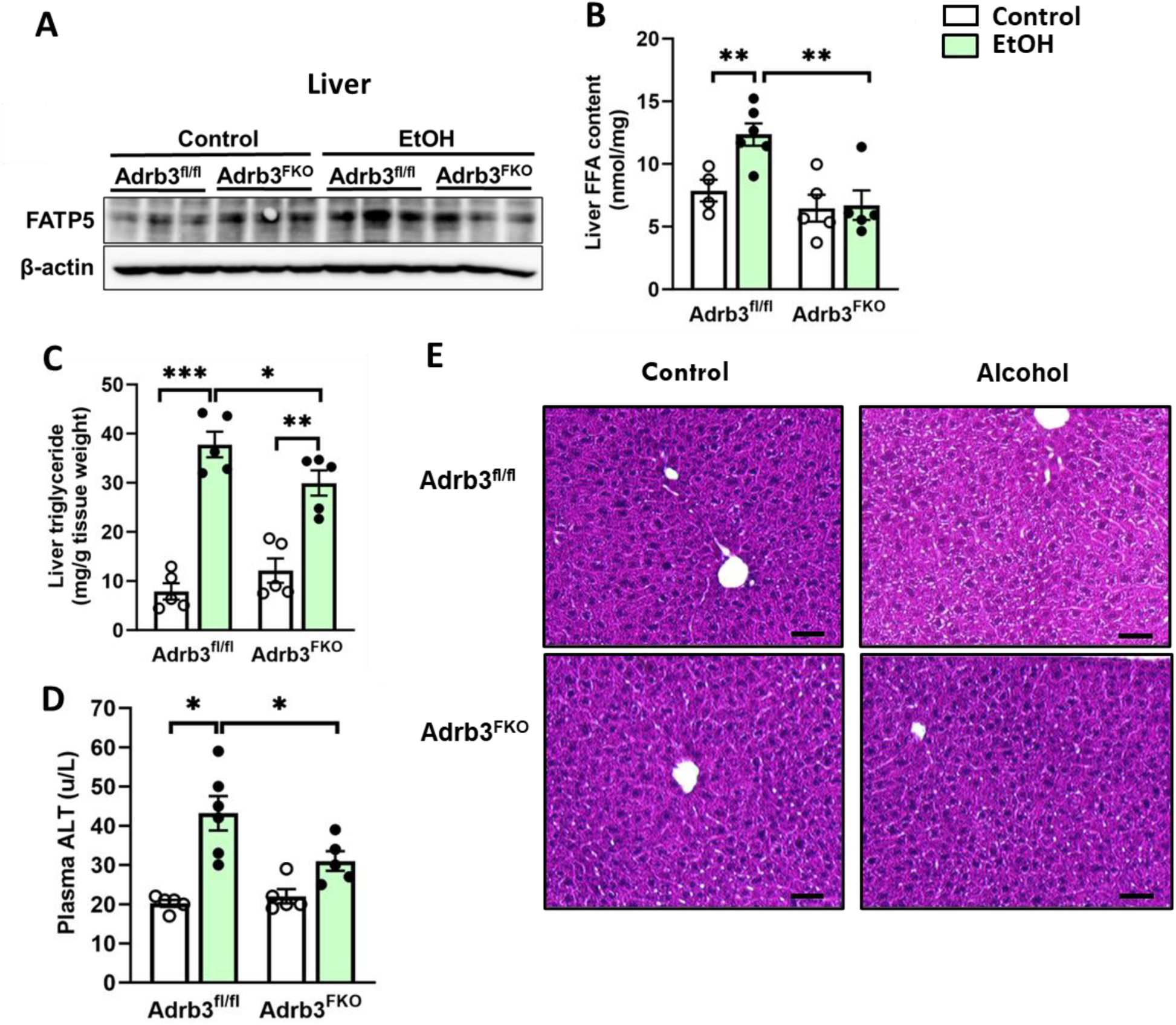
Ablation of Adrb3 in adipose tissues attenuates acute alcohol-induced hepatic steatosis and liver damage. Chow-fed Adrb3^fl/fl^ and Adrb3^FKO^ mice were given a single alcohol binge. (A) Liver FATP5 protein expression. (B) Liver FFA. (C) Liver triglyceride. (D) Plasma ALT. (E) H&E staining of liver tissue sections. Scale bar: 50 µm. n=5-6. Data are expressed as the means ± SEM. *P < .05, **P < .01, *** P < .001.

In addition to enhanced fatty acid uptake, other factors, including elevated endogenous lipogenesis, downregulated fatty acid β-oxidation, and reduced lipid export also contribute to ALD development.^22^ Hepatic expression of genes involved in lipogenesis was examined and we found that acute alcohol treatment suppressed hepatic expression of SREBP1c and SCD1 in both Adrb3^fl/fl^ and Adrb3^FKO^ mice (**Figure S5A**). Hepatic FASN mRNA expression was slightly but not significantly increased in both genotypes. (**Figure S5C**). Interestingly, binge drinking caused significant reductions in ACC and DGAT1 mRNA expression in the liver of Adrb3^FKO^ mice compared to that of Adrb3^fl/fl^ mice (**Figure S5A and S5B**). Either acute alcohol or adipocyte Adrb3 deficiency affected hepatic DGAT2 expression (**Figure S5B**). Regardless of the genotype, hepatic levels of genes critical for β-oxidation (PPARα, CPT1α and Acox1) and lipoprotein formation (apolipoprotein B) were comparably suppressed after acute alcohol treatment (**Figure S5C-D**). Taken together, these results indicated that adipose Adrb3 deficiency could protect against acute alcohol-induced fatty liver and liver damage mainly via attenuated hepatic fatty acid uptake.

### Adipocyte Adrb3 deficiency is associated with elevated plasma leptin and enhanced hepatic pAKT expression following binge alcohol exposure

Activation of ADRB3 signaling has been shown to suppress leptin expression.^23^ Therefore, we sought out to determine whether adipocyte Adrb3 deficiency could enhance leptin levels in the content of alcohol drinking. We found that circulating leptin levels were not affected by adipocyte Adrb3 ablation in control-gavaged mice, but significantly elevated following binge drinking (**Figure 8A**). This increase in plasma leptin was correlated with elevated hepatic leptin receptors expression in acute alcohol-exposed Adrb3^FKO^ mice (**Figure 8B-C**). Accumulating evidence suggests that alcohol consumption leads to hepatic insulin resistance.^8^ Supporting this, we observed a significant reduction of pAKT expression in Adrb3^fl/fl^ mice after acute alcohol treatment. Interestingly, hepatic pAKT expression was maintained in Adrb3^FKO^ mice (**Figure 8D**).

**Figure 8.**
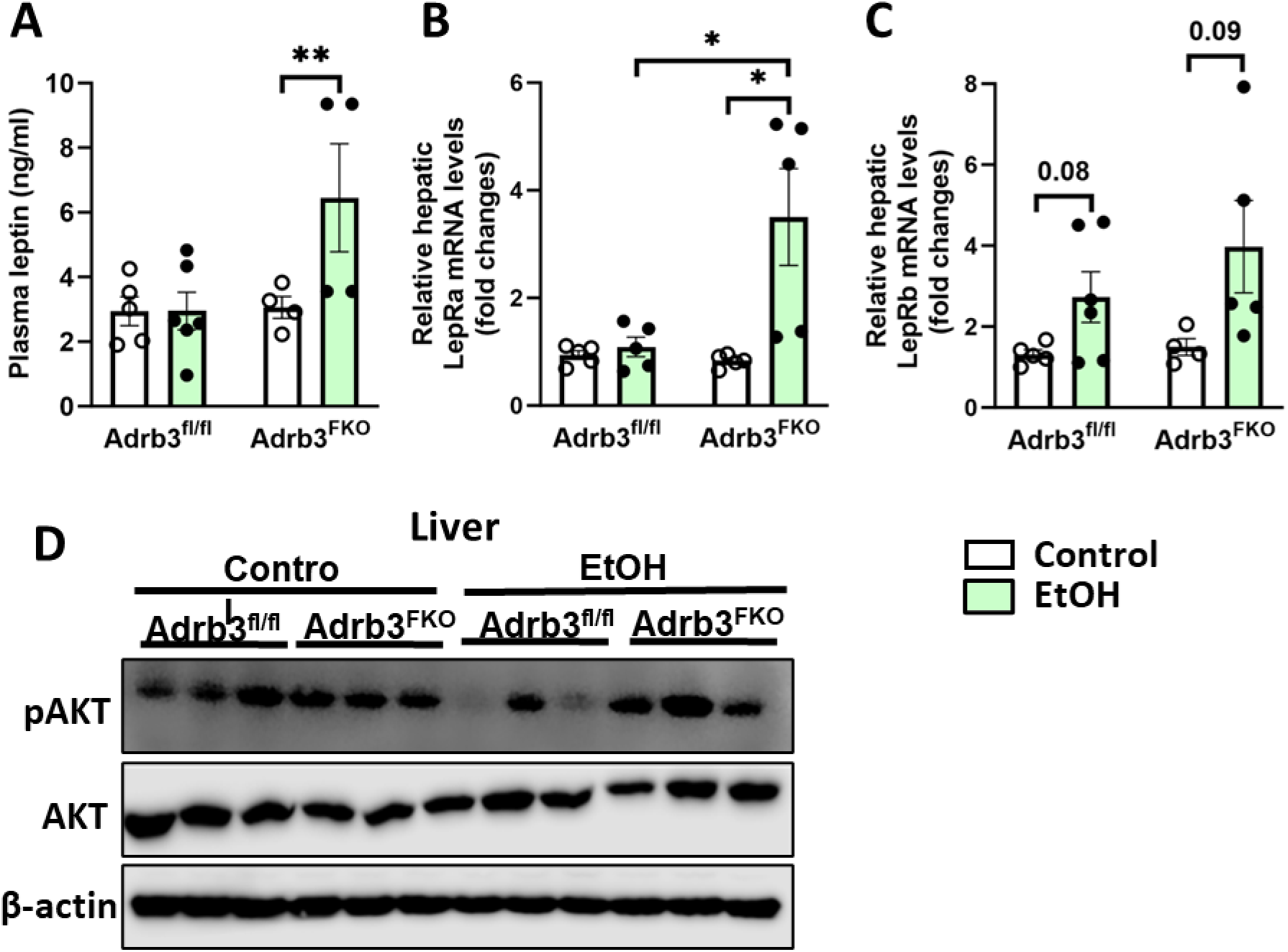
Mice lacking adipocyte Adrb3 show elevated plasma leptin and enhanced hepatic pAKT expression following binge drinking. Chow-fed Adrb3^fl/fl^ and Adrb3^FKO^ mice were exposed to acute alcohol treatment. (A) Plasma leptin level. (B-C) RT-qPCR analysis of (B) LepRa and (C) LepRb in the liver. (D) Liver pAKT protein expression. n=4-6. Data are expressed as the means ± SEM. *P < .05, **P < .01.

## Discussion

Acute alcohol drinking has become a serious health problem worldwide. Over 90% of US adults who drink excessively report binge drinking.^2,24,25^ Despite the highlighted role of adipose tissue lipolysis in the development of alcoholic liver injury,^7,9^ the precise mechanisms by which acute alcohol exposure enhances adipose tissue lipolysis are not fully understood.^25^ Catecholamines are key players in triglyceride hydrolysis and FFA release from lipid droplets by interacting with β-ARs and activating a series of kinase-mediated phosphorylation cascades. Elevated circulating NE and epinephrine levels were observed in NIAAA model-fed mice.^39^ In contrast, Zhong et al. reported that chronic alcohol feeding did not affect plasma NE or epinephrine contents.^8^ Nevertheless, it has been shown that adipocyte lipolysis is more likely to be stimulated by the high concentrations of NE that is found in the immediate vicinity of the terminal adrenergic nerve fibers than by catecholamines derived from the circulation.^15^ In agreement, adrenal demedullation did not protect rats from acute alcohol consumption-induced liver lipid accumulation, indicating that adrenal glands secreted NE or epinephrine does not affect binge drinking-associated lipolysis and fatty liver development.^26^ In the current study, we found that acute alcohol drinking caused significant increases in local adipose tissue NE, which could activates β-ARs-mediated lipolytic pathway, contributing to hepatic fat accumulation.

The important role of alcohol-induced SNS activation in adipose tissues and subsequent fatty liver development has been reported in mice following alcohol overconsumption. Specifically, Song et al. found that surgical denervation of both sympathetic and sensory nerves attenuates chronic alcohol-induced elevations in plasma FFA and hepatic fat contents.^9^ It is worth noting that sensory nerves may also play a role in mediating adipose tissue lipolysis via sending signals to certain brain regions where integrated information can stimulate peripheral lipolysis.^27^ Similarly, Zhou et al. reported that mice lacking peripheral tyrosine hydroxylase (TH), a marker of both sympathetic and sensory neurons, exhibit reductions in lipolysis and hepatic steatosis when they were challenged by repeated binge drinking.^12,27^ When mice were housed at thermoneutrality, Ibras et al. reported that 6-OHDA-mediated SNS disruption led to reduced hepatic triglyceride levels when mice were exposed to binge drinking. However, this study did not investigate the lipolytic function of adipose tissues. Using a single binge drinking model, we observed that both systemic and local denervation of sympathetic nerves protected mice from acute alcohol-induced increases in adipose tissue pHSL expression and plasma FFA as well as development of alcoholic fatty liver disease. Collectively, our findings show that blocking NE-mediated adipocyte lipolysis protects mice from acute alcohol-induced adipose tissue hyper-lipolysis, hepatic steatosis, and liver damage.

Controversial findings have been reported regarding the combined effect of β-AR agonist and alcohol feeding on adipose tissue lipolysis. When rats were fed alcohol chronically for 4 weeks, an intraperitoneal injection of CL316,243 led to sustained reductions in plasma FFA for 60 min.^29^ In addition, decreased β-AR-stimulated lipolysis was observed in cultured adipocytes isolated from rat exposed to chronic alcohol feeding.^30^ Conversely, Song et al. found that 7-day CL316,243 administration further increased plasma FFA levels in chronic alcohol-fed mice, which was accompanied by dramatically increased hepatic triglyceride.^9^ Consistently, we observed a synergistic effect of single binge drinking and CL316,243 administration on fatty liver development despite similar circulating FFA levels between EtOH+Saline and EtOH+CL316, 243-treated mice 5.5 hours following alcohol oral gavage. The discrepancies among these studies could be due to the differences in rodent models, alcohol drinking patterns, and/or durations of ADRB3 agonist treatment.

Among the three β-ARs, Adrb3 is highly expressed in adipose tissues.^6^ The finding that Adrb3 antagonist-treated mice are protected from chronic alcohol-induced elevations in plasma glycerol and hepatic triglycerides^9^ led us to determine the fat-specific role of Adrb3 in acute alcohol-associated adipocyte lipolysis and liver damage. Intriguingly, we found that relative to acute alcohol-fed Adrb3^fl/fl^ mice, Adrb3^FKO^ mice exhibited significant reductions in plasma FFA, liver lipid and circulating ALT. The partial protection from ALD in these mice could result from decreased Adra2 expression in adipose tissues since Adra2 suppresses lipolysis by interacting with Gi proteins. It would be interesting to investigate in the future whether activation of Adra2 in Adrb3^FKO^ mice could exert more protective effects on ALD.

Studies in humans have shown that serum leptin concentration was reduced by either chronic or acute alcohol consumption.^31–34^ Tan et al. reported that chronic alcohol exposure downregulates leptin expression in WAT and reduces plasma leptin levels in mice.^34^ By normalizing circulating leptin levels, leptin administration rescues alcoholic fatty liver development.^34^ Unlike chronic alcohol feeding, binge drinking did not alter plasma leptin as shown in Figure 8A. Interestingly, adipocyte Adrb3 deficiency was associated with elevated circulating leptin in acute alcohol-treated mice. This observation is consistent with the concept that adrb3 signaling suppresses leptin secretion.^23^ Together with the increased hepatic expression of leptin receptors and the important role of leptin deficiency in the pathogenesis ALD, enhanced leptin signaling in the liver could partially contribute to attenuated liver injury and enhanced insulin signaling observed in adipocyte Adrb3 ablated mice.^34–36^

With the increase prevalence of binge alcohol drinking, especially among college students and other populations, binge drinking has emerged as a critical public health concern and a substantial risk factor for the development of severe ALD. Therefore, much research attention is afforded to binge alcohol drinking. Here we report that acute alcohol intake is associated with increased NE in WAT. Systemic and local ablation of sympathetic nerve attenuated binge drinking-induced adipose lipolysis and fatty liver disease. We also provided evidence supporting an important role of adipocyte Adrb3 in mediating acute alcohol-induced hyper-lipolysis in isolated fat pads. Another key finding of our study was that Adrb3 deficiency in adipose tissues partially but significantly attenuated acute alcohol-induced increases in plasma FFA and liver fat accumulation.

## Materials and Methods

### Animal care

Animals were housed in a pathogen free barrier facility with a 12h light-dark cycle (6:00 a.m.-6:00 p.m.). They had free access to food (standard chow diet, 2916 Global Diet; Harlan Teklad) and water unless specified otherwise. Experiments were performed according to protocols reviewed and approved by the Institutional Animal Care and Use Committee of the University of Texas at Dallas (UTD).

### Single oral gavage of alcohol

Male C57BL/6J mice (10-12 weeks old) were given an oral gavage of 5 g/kg BW of alcohol (31.5%, vol/vol, referred to as EtOH) or maltose dextrin (45%, wt/vol, referred to as Control). Six hours later, mice were anesthetized for blood, liver, gWAT, and iWAT collection. Tissues were quickly removed, snap-frozen in liquid nitrogen and stored at -80 °C.

### Systemic 6-OHDA treatment

To ablate sympathetic nerves, intraperitoneal injections of 6-hydroxydopamine (referred to as 6-OHDA, H4387, Sigma-Aldrich) at 1mg/kg BW were performed in male C57BL/6 mice (Jackson Laboratory) (10-12 weeks old). 6-OHDA was prepared in 0.1% ascorbic acid (Thermo Scientific™, Waltham, MA, USA) and 0.9% sterile NaCl (referred to as saline) (Hospira). Injections were performed for three consecutive days.^37^ Then on the fourth day, both 6-OHDA injected and saline injected mice were oral gavaged with 5 g/kg BW alcohol or 45% of maltose dextrin in the early morning. Six hours later, mice were anesthetized for blood and tissue collection.

### Chemical denervation of gonadal fat pads

Chemical ablation of sympathetic nerves in gonadal WAT (gWAT) was performed as previously described.^37^ Briefly, male C57BL/6J mice (10-12 weeks old) were anesthetized and the incision area was cleaned by hair removal and 75% alcohol gauze wipe treatment. An incision of appropriate length along the mouse abdominal midline was made so that bilateral gWAT could be explored during surgery. Sterile saline was applied to keep the tissue moist. 10 loci across the gWAT pad were injected with 2 μl of 6-OHDA (8 mg/ml). After the surgery, the tissue and incision were washed with saline before closing the muscle tissue and skin of the incision with silk sutures and wound clips, respectively. Mice were intraperitoneally injected with ketoprofen (5mg/kg BW) every 24 hours for 3 days to prevent infection. Mice were subjected to alcohol or maltose dextrin binge after 2 weeks of recovery.^37^

### Histological analysis

Adipose tissues and liver samples were fixed in 10% formalin and processed for paraffin embedding at the Histological Core Facility at UTD. Paraffin tissue sections were cut at 5 μm and 10 μm for adipose tissue and liver samples, respectively. Hematoxylin and eosin (H&E) staining was conducted and the whole mount images were obtained using a Zeiss microscope.

### Generation of Adrb3^fl/fl^ mice

Conditional mouse model for the Adrb3 gene was developed using Alt-R® CRISPR-Cas9 System from Integrated DNA Technologies in Dr. Joel Elmquist laboratory at the University of Texas Southwestern Medical Center (UTSW). Briefly, two synthetic sgRNA targeting introns 1 and 3 (crRNA 1: 5’-GGG GAC GGA TCT CAT CAG CTG GG-3’; crRNA 2: 5’-GGG TGC AGG ATG GAC CCT ATG GG -3’) were synthetized and co-injected with the tracrRNA, two Ultramer® DNA Oligonucleotides containing the LoxP sequence (ATA ACT TCG TAT AGC ATA CAT TAT ACG AAG TTA T) and ∼60 bp of homology arm, and Alt-R S.p Cas9 Nuclease 3NLS, in C57BL6 ES cells by the UTSW Transgenic Core (Figure S2A-C). Guides were selected using the CRISPR Design Tool (http://tools.genome-engineering.org). Mice were screened and the founder backcrossed with a C57BL6 mouse to ensure germ line transmission. The heterozygous (Adrb3^fl/+^) F1 pups were then crossed together to generate homozygous (Adrb3^fl/fl^) mice (Figure S2D), which were maintained until bred with an Adiponectin (Adipoq)-Cre mouse (028020; The Jakson Laboratory) to generate Adrb3^FKO^ at UDT.

### CL316,243 treatment

30 min after a single binge drinking, male C57BL/6J mice were intraperitoneally injected with 0.1mg/kg BW of CL316,243 or saline. Blood was collected at 0, 1, 2, 4, and 5.5 hours following CL316,243 administration. Livers were collected for the determination of liver triglyceride content at 5.5 hours after CL316,243 injection.

Chow-fed Adrb3^fl/fl^ and Adrb3^FKO^ male mice (8-10 weeks old) were treated with 0.1mg/kg BW of CL316,243 via intraperitoneal injection. Blood was collected at different time points (0, 0.5, 1, 2, and 4 hours) after CL316,243 administration to determine blood glucose levels and plasma FFA contents.

### Measurement of liver triglyceride contents

Frozen liver tissues (60–80 mg) were thawed, minced, and weighed in glass tubes. Lipids were extracted in 3 mL of 2:1 chloroform: methanol at room temperature overnight. After centrifugation, lipid extract was transferred to a clean glass tube, and dilute H_2_SO_4_ (0.05%) was added to separate the phases by vortex and centrifugation. The aqueous upper phase was removed, and a 100 µL aliquot of the bottom phase was transferred to a new glass tube and dried down under N_2_. Then, 1 mL of 1% Triton X-100 in chloroform was added. After the evaporation of the solvent, 500 µL of deionized water was added to each tube and vortexed until the solution was clear. Triglyceride standards (were prepared by adding 1% Triton X-100 in chloroform, evaporating, and dissolving in deionized water. Triglyceride contents in liver samples were quantified using the Infinity Triglycerides Reagent (Thermo Scientific™, Waltham, MA, USA).

### Western blotting

gWAT and iWAT samples (pooled samples from the same genotype and treatment, n=2– 3) were homogenized in lysis buffer containing 50 mM Tris-HCl (pH 8.0), 0.25 mol/L NaCl 5 mM EDTA, 1% Triton X-100 (v/v), 1% NP-40, protease inhibitor (P8340, Sigma, St. Louis, MO, USA) and phosphatase inhibitor cocktails (P5726 and P0044, Sigma, St. Louis, MO, USA).^38^ Liver samples were homogenized in RIPA buffer containing 25 mM Tris-HCl (pH 8.0), 150 mM NaCl, 1% NP-40, 1% sodium deoxycholate, 0.1% SDS (pH 7.6), protease inhibitor (P8340, Sigma, St. Louis, MO, USA) and phosphatase inhibitor cocktails (P5726 and P0044, Sigma, St. Louis, MO, USA). Protein concentrations were determined by BCA kit (Pierce, Thermo Scientific™, Waltham, MA, USA). 10-20 µg tissue lysates were separated by 8% gel in SDS-PAGE and transferred to nitrocellulose membranes (Trans-Blot, Bio-Rad, Hercules, CA, USA). Membranes were blocked with 5% nonfat dried milk or 3% BSA (for phosphorylated proteins) for 1 h at room temperature and then incubated with primary antibodies overnight at 4°C in 5% nonfat dried milk or 3% BSA (for phosphorylated proteins). Primary antibodies were purchased from Cell Signaling Technology, phospho-HSL (Ser473) (#9271), total HSL (#9315), FATP5 (#NBP2-37412), GAPDH (#2118S) and β-Actin (#4970S). Membranes were washed with TBS containing 0.1% (vol/vol) Tween 20, and incubated with 1:10,000 dilution of goat anti-rabbit horseradish peroxidase antibody (Jackson Immuno Research, West Grove, PA, USA) for 1 h at room temperature. Blots were visualized with enhanced chemiluminescence (Bio-Rad, Hercules, CA, USA).

### Plasma parameters

Mice were anesthetized and blood was collected in EDTA-coated tubes. Plasma was separated by centrifugation at 8,000 g for 15 min and stored at -80°C. Plasma alanine transaminase (ALT) was measured either using the Vitros 250 Chemistry Analyzer (Ortho Clinical Diagnostics) by Metabolic Phenotyping Core at UTSW or using a commercial kit (Teco Diagnostics). Plasma FFA contents were measured using NEFA-WAKO Assay Kit (Wako) following the manufacturer’s instructions.

### Measurement of hepatic free fatty acid (FFA) concentration

Liver samples were homogenized in 200 µL chloroform/Triton X-100 (1% Triton X-100 in pure chloroform). Homogenized samples were incubated on ice for 10-30 minutes and centrifuged for 5-10 minutes at top speed in a microcentrifuge. Lower organic phase was collected and air dried at 50°C in a fume hood to remove chloroform. Dried lipids were dissolved in 200 µL of assay buffer by vertexing extensively for 5 minutes. Hepatic FFA contents were measured using a Free Fatty Acid Assay Kit (Abcam) following the manufacturer’s instructions.

### Ex vivo lipolysis assay

Two hours after a single binge of alcohol or maltose dextrin, gWAT and iWAT of Adrb3^fl/fl^ and Adrb3^FKO^ mice were surgically removed, washed in DPBS and incubated in prewarmed (37 °C) Dulbecco’s modified Eagle’s medium (DMEM, 1 g/L glucose; GIBCO, Life Technologies, Carlsbad, CA) until it is ready for use. Tissue pieces (∼20 mg) were cut and incubated in 200 μl DMEM containing 2% BSA (FA-free) (Sigma-Aldrich) in the presence of 5 μM triacsin C (Sigma-Aldrich) in 96-well plates at 37 °C, 5% CO_2_, and 95% humidified atmosphere for 1 hour. Then culture medium was removed for FFA measurement. Another 200 μl fresh DMEM medium containing 2% FA-free BSA and triacsin C (5 μM) was added to fat explant and collected after 1 h incubation for the determination of FFA content. The fat explants were transferred into 1 ml extraction solution (chloroform/methanol (2:1, v/v) in 1% glacial acetic acid) and incubated for 120 minutes at 37 °C under vigorous shaking on a thermomixer. Fat explants were transferred into 500 µl lysis solution (NaOH/SDS (0.3 N/0.1%) and incubated overnight at 55 °C under vigorous shaking on a thermomixer. The protein content of the fat explant lysates was determined using BCA kit (Pierce, Thermo Scientific™, Waltham, MA, USA). The FFA in the incubation media were determined using a NEFA kit (Wako chemicals). The result was normalized to protein concentration of the tissues used for the assay.

### RT-qPCR

Total RNAs from liver, gWAT, and iWAT were extracted using RNA Stat60 (Tel-Test). Complementary DNA was synthesized using the iScript Advanced cDNA Synthesis Kit for RT-qPCR (Bio-Rad). qPCR was performed using a Bio-Rad sequence detection system (Bio-Rad). 18s and GAPDH were used as internal controls for liver and fat tissues respectively. The relative amounts of all mRNAs were calculated using the ΔΔCT assay. The mouse primer sequences are shown in Table 1 and were obtained from Integrated DNA Technologies (IDT).

**Table 1.**
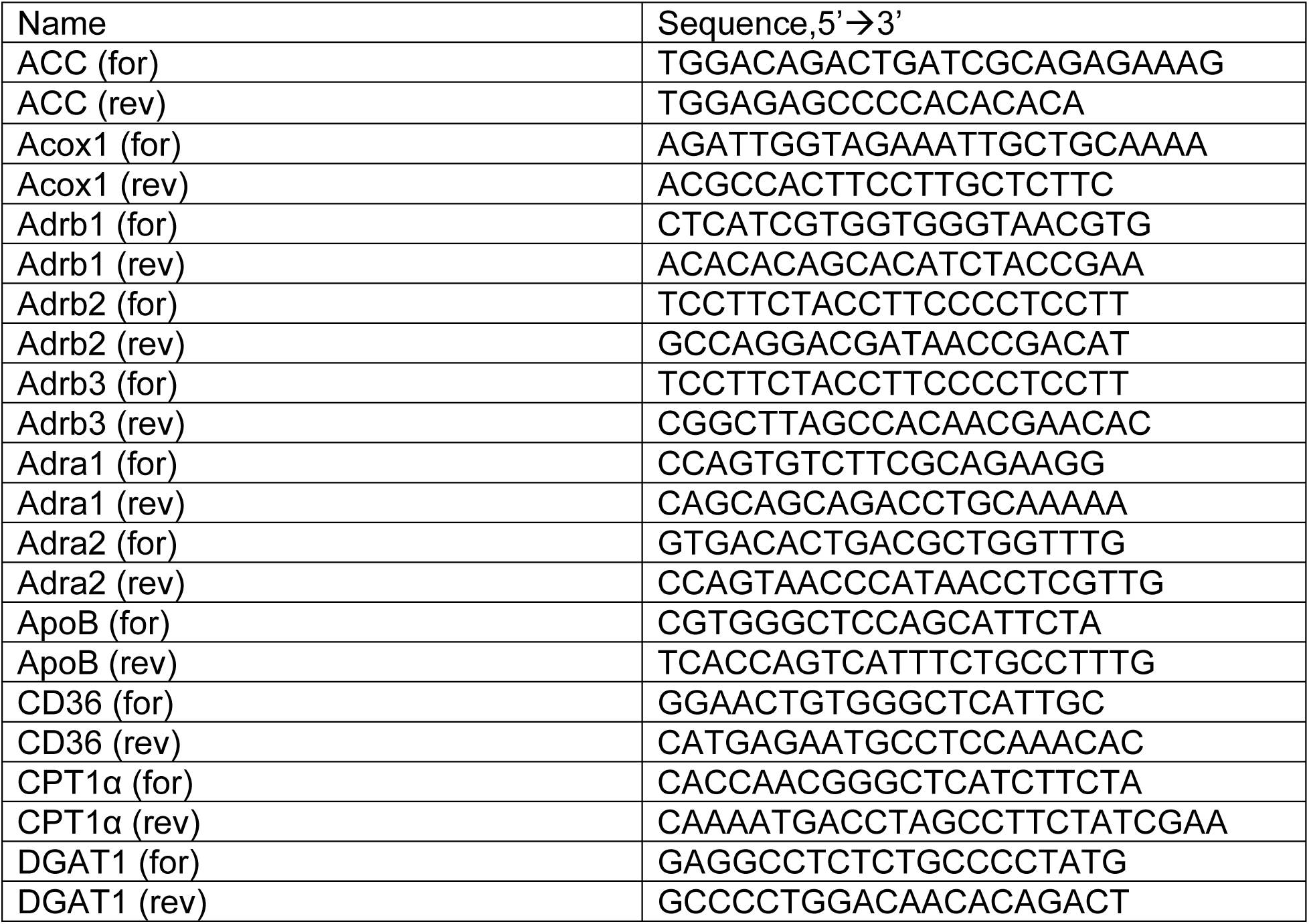
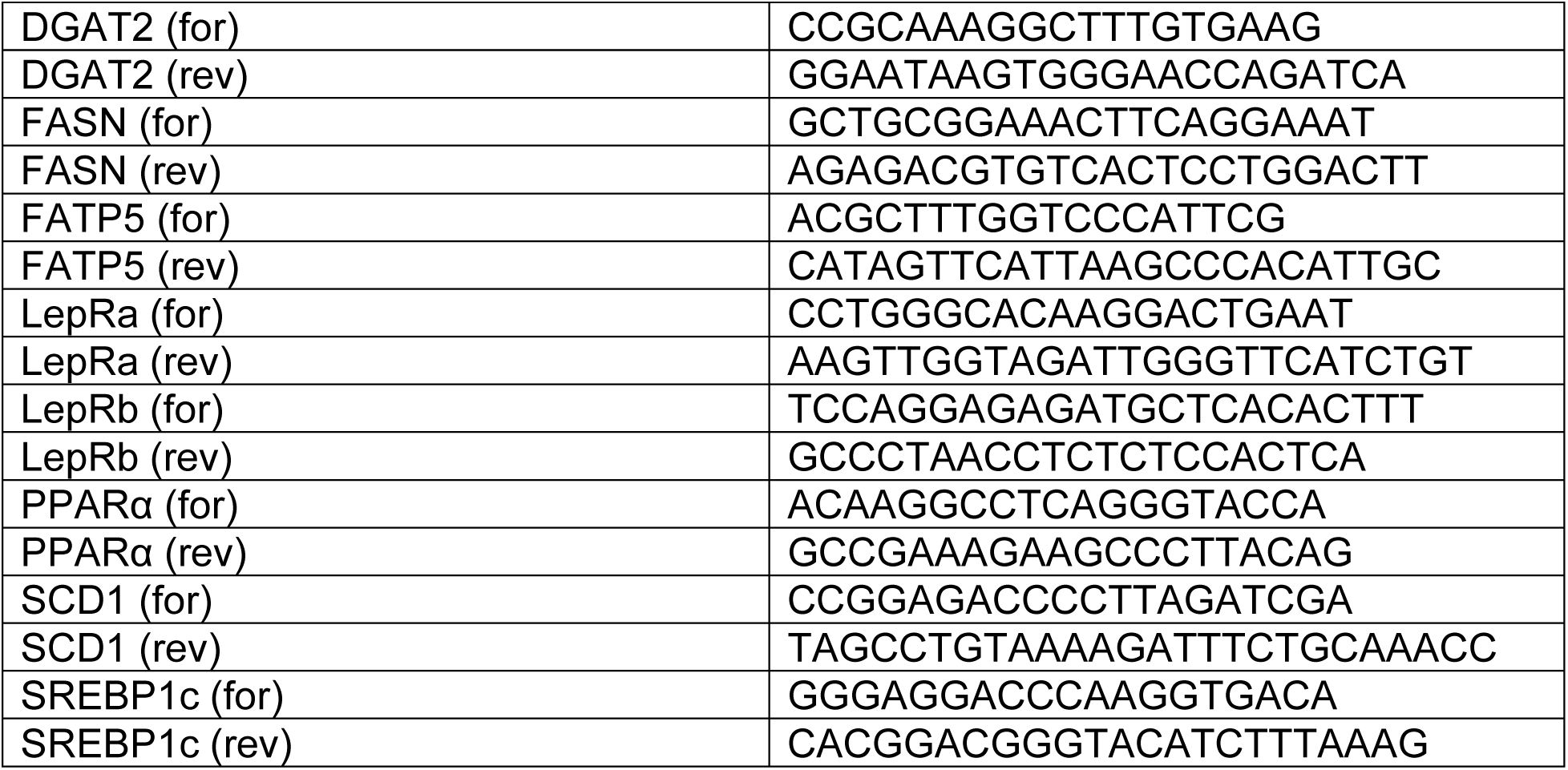
List of Primers.

### Statistical analysis

Data are expressed as means ± SEM. Statistical analysis was performed using Student’s t-test for experiments comparing two groups. For studies with three or more groups, a one-way ANOVA was used (GraphPad Prism, Boston, FL, USA). p<0.05 is considered significant.

## Abbreviations used in the paper

ALD: alcohol-associated liver disease
β-ARs: β adrenergic receptors
Adrb3: β3 adrenergic receptor
NE: norepinephrine
SNS: sympathetic nervous system
pHSL: phosphorylated hormone-sensitive lipase
FFA: free fatty acid
6-OHDA: 6-hydroxydopamine
gWAT: gonadal white adipose tissue
iWAT: inguinal white adipose tissue
WAT: white adipose tissue.

---

**Supplemental figure 1.**
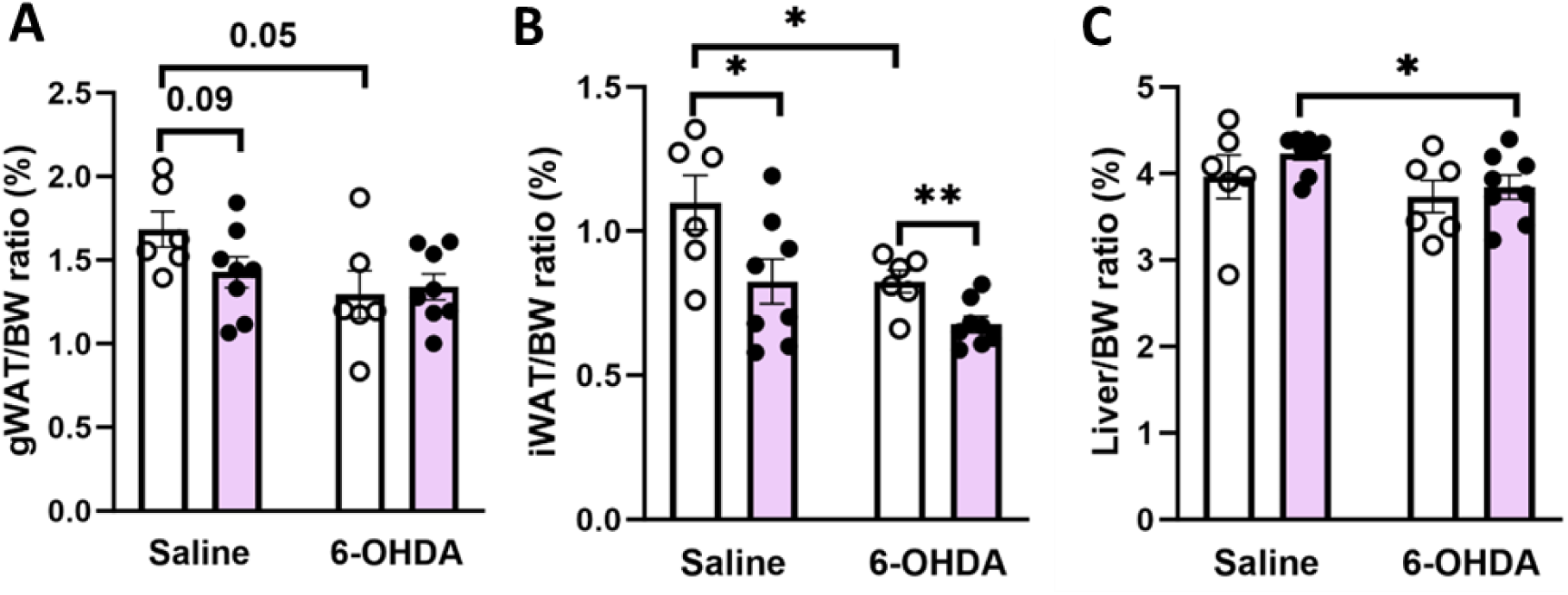
Tissue to body weight (BW) ratios in binge drinking-exposed mice following systemic 6-OHDA treatment. Mice were intraperitoneally injected with 6-OHDA once per day for 3 days. On the fourth day, mice were subjected to an oral gavage of alcohol (EtOH) or isocaloric maltose dextrin (Control). (A) gWAT/BW ratio (%). (B) iWAT/BW ratio (%). (C) Liver/BW ratio. Data are expressed as the means ± SEM. n=5-8. *P < .05, **P < .01. 6-OHDA, 6-Hyroxydopamine.

**Supplemental figure 2.**
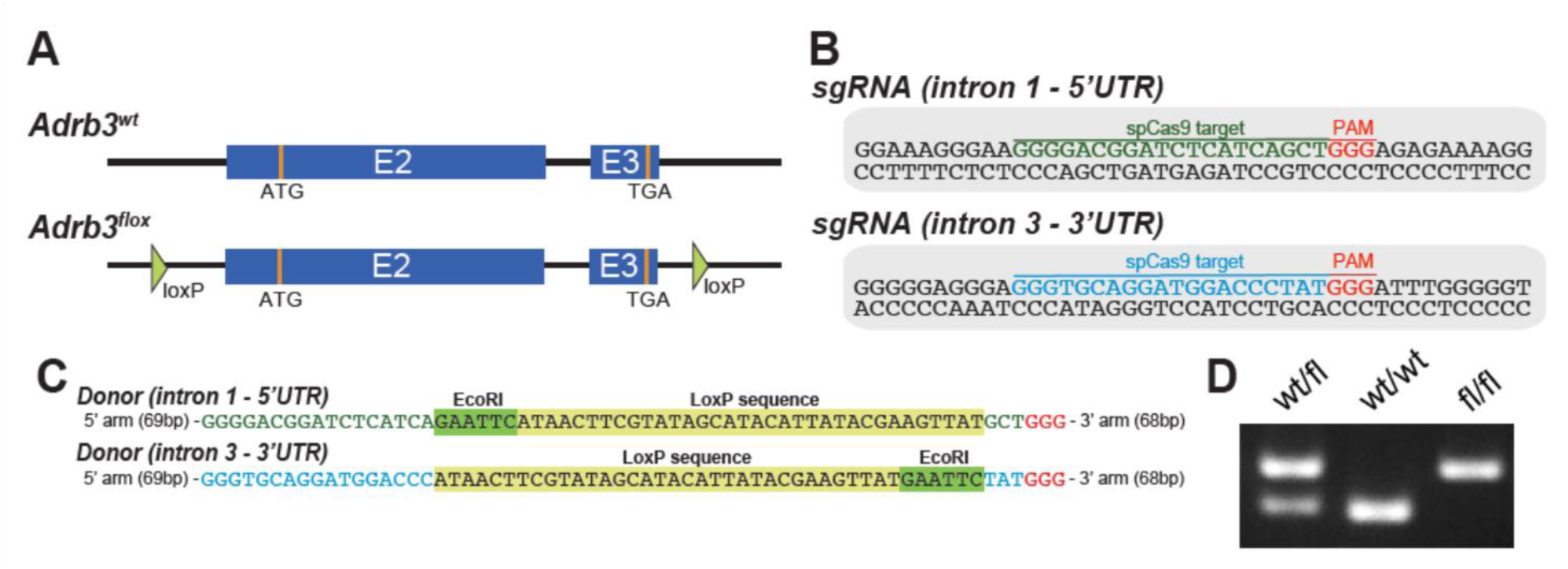
Generation of the Adrb3^fl/fl^ mouse model. Adrb3^fl/fl^ mice on a C57BL/6 background were generated using CRISPR-Cas9 technology. (A) Gene targeting strategy showing the generation of mice with floxed Adrb3 allele. (B) sgRNA sequence. (C) TraceRNA containing the LoxP sequence. (D) PCR results of wild type (wt) or floxed allele (fl).

**Supplemental figure 3.**
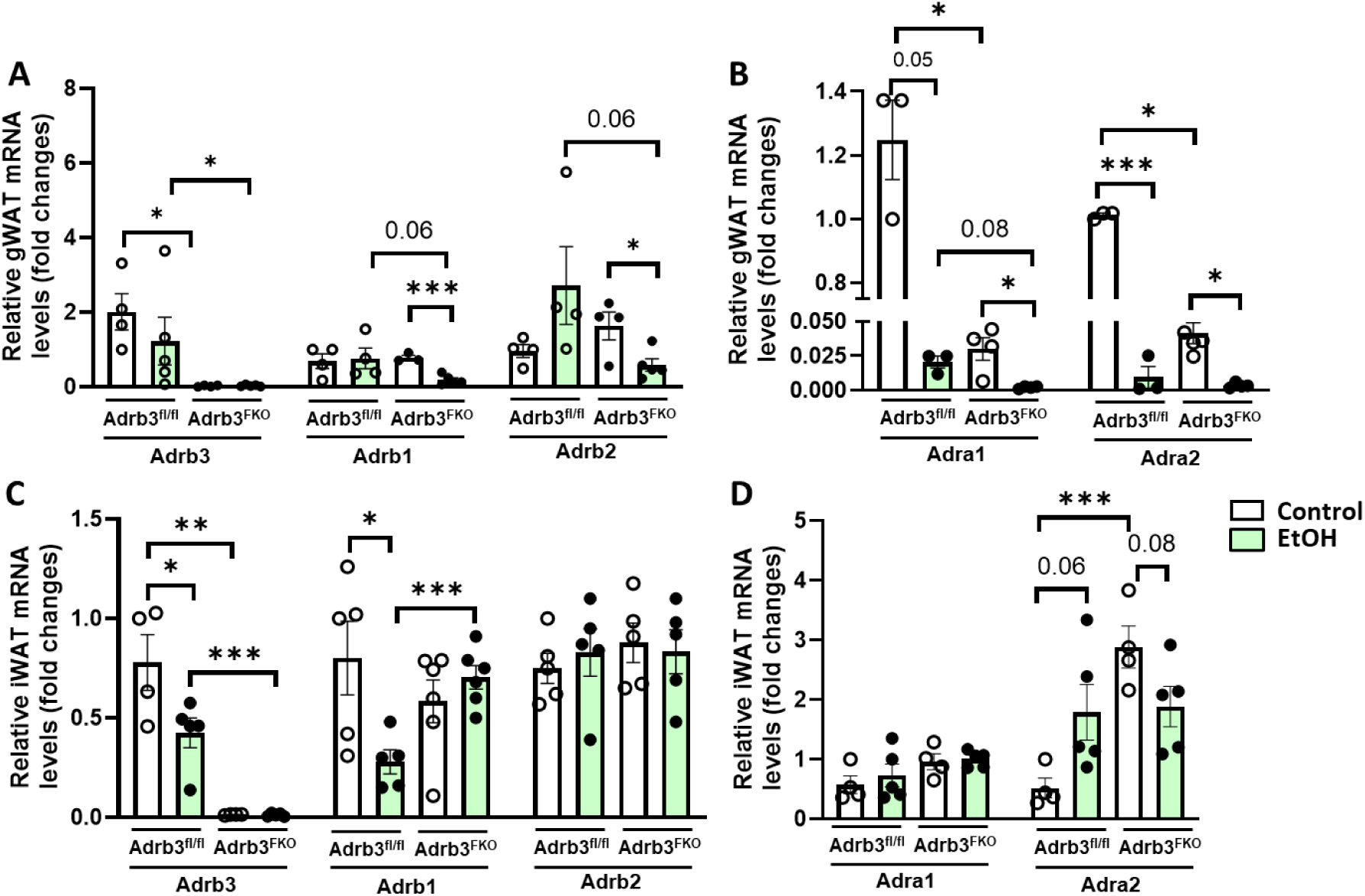
Expression of adrenergic receptors (ARs) in fat pads of mice following binge drinking. RT-qPCR analysis was performed to detect expression of β-ARs in (A) gWAT and (C) iWAT. Expression of Adra1 and Adra2 in (B) gWAT and (D) iWAT. Data are expressed as the means ± SEM. n=3-5. *P < .05, **P < .01, *** P < .001.

**Supplemental figure 4.**
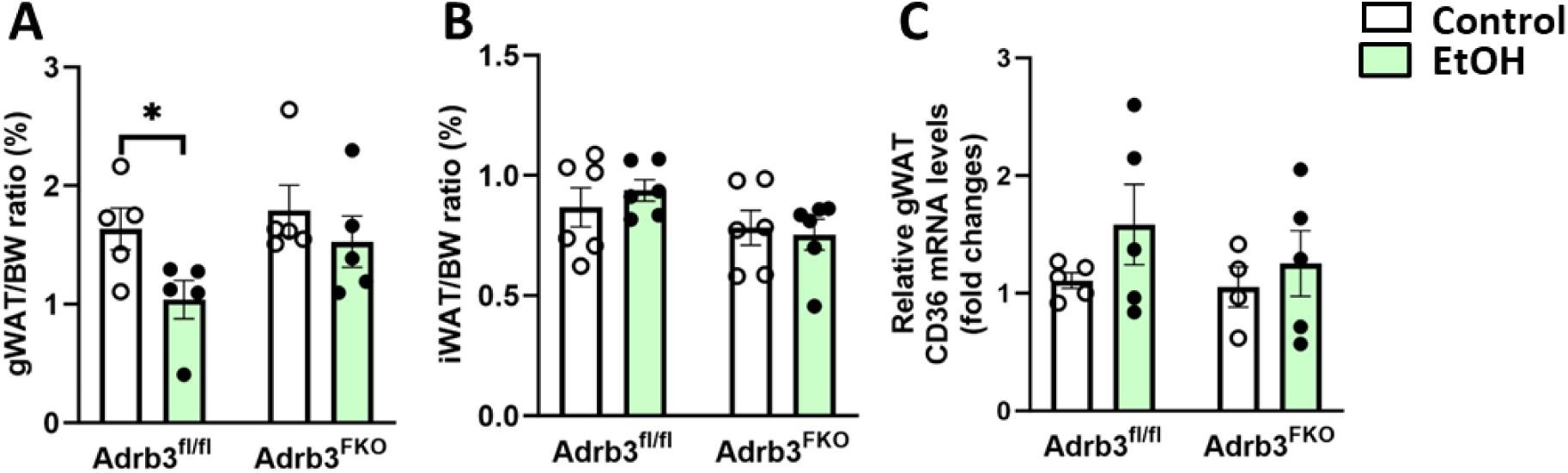
Binge drinking reduces gWAT/BW ratio in Adrb^fl/fl^ mice. (A) gWAT/BW ratio (%). (B) iWAT/BW ratio (%). (C) RT-qPCR analysis of hepatic CD36 expression. Data are expressed as the means ± SEM. n=4-6. *P < .05.

**Supplemental figure 5.**
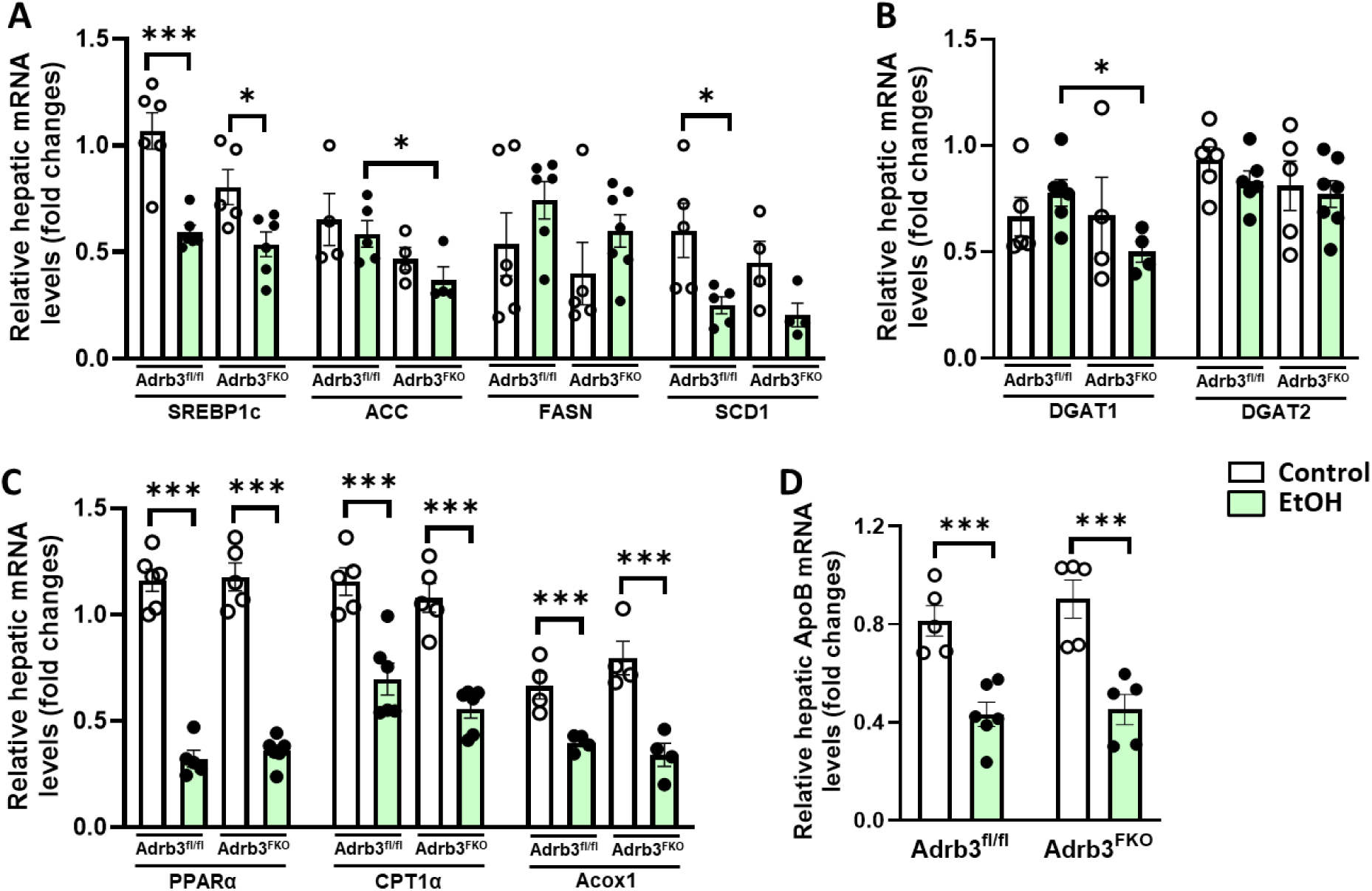
Deletion of the Adrb3 gene in adipose tissues does not affect the expression of genes involved in de novo lipogenesis, fatty acid beta-oxidation, and lipid export. Chow-fed Adrb3^fl/fl^ and Adrb3^FKO^ mice were given a single alcohol binge. Mice were killed 6 hours after alcohol gavage. RT-qPCR was performed using liver samples for the detection of (A) SREBP1c, ACC, FASN and SCD1; (B) DGAT1 and DGAT2; (C) PPARα, CPT1α and Acox1; (D) ApoB. n=4-6. Data are expressed as the means ± SEM. n=4-6. *P < .05, *** P < .001.

